# Generation of Recombinant Mammalian Selenoproteins through Genetic Code Expansion with Photocaged Selenocysteine

**DOI:** 10.1101/759662

**Authors:** Jennifer C. Peeler, Rachel E. Kelemen, Masahiro Abo, Laura C. Edinger, Jingjia Chen, Abhishek Chatterjee, Eranthie Weerapana

## Abstract

Selenoproteins contain the amino acid selenocysteine and are found in all domains of life. The functions of many selenoproteins are poorly understood, partly due to difficulties in producing recombinant selenoproteins for cell-biological evaluation. Endogenous mammalian selenoproteins are produced through a non-canonical translation mechanism requiring suppression of the UGA stop codon, and a selenocysteine insertion sequence (SECIS) element in the 3’ untranslated region of the mRNA. Here, recombinant selenoproteins are generated in mammalian cells through genetic code expansion, circumventing the requirement for the SECIS element, and selenium availability. An engineered orthogonal *E. coli* leucyl-tRNA synthetase/tRNA pair is used to incorporate a photocaged selenocysteine (DMNB-Sec) at the UAG amber stop codon. Recombinantly expressed selenoproteins can be photoactivated in living cells with spatial and temporal control. Using this approach, the native selenoprotein methionine-*R*-sulfoxide reductase 1 is generated and activated in mammalian cells. The ability to site-specifically introduce selenocysteine directly in mammalian cells, and temporally modulate selenoprotein activity, will aid in the characterization of mammalian selenoprotein function.

---

Selenoproteins contain the amino acid selenocysteine (Sec), which is a cysteine (Cys) cognate with selenium (Se) in place of sulfur (S). In the majority of characterized selenoproteins, such as glutathione peroxidase and thioredoxin reductases, Sec performs redox-catalytic functions^1^. Selenoproteins are found throughout all three domains of life, with 25 selenoproteins in humans.^2^ However, some species express Cys-containing homologs of selenoproteins, suggesting that Cys can nominally perform the same function as Sec. The stringent requirement for Sec in certain organisms is attributed to the reversible nature of Sec oxidation, which confers protection in contrast to irreversible Cys oxidation.^3^ Replacement of Sec with Cys in glutathione peroxidase 4 (GPX4) renders neurons susceptible to ferroptotic cell death due to overoxidation and inactivation of GPX4-Cys.^4^ This observation demonstrates a potential advantage conferred by the energetically expensive production of selenoproteins.

Sec incorporation deviates from canonical protein translation, requiring suppression of the UGA stop codon. In eukaryotes, Sec biosynthesis occurs directly on the suppressor tRNA (tRNA^[Ser]Sec^). Specifically, tRNA^[Ser]Sec^ is aminoacylated with serine by seryl-tRNA synthetase (SerS), followed by phosphorylation by phosphoseryl-tRNA kinase (PSTK), and subsequent Se incorporation by Sec synthase (SecS), to generate Sec-tRNA^[Ser]Sec^ (Figure 1a).^5,6^ Suppression of UGA by Sec-tRNA^[Ser]Sec^ requires a Sec insertion sequence (SECIS) element in the selenoprotein mRNA. The SECIS appears in the 3’ UTR of the selenoprotein mRNA, and binding of the SECIS binding protein 2 (SBP2) and Sec-specific translation elongation factor (EFSec) are required for successful Sec insertion (Figure 1a).^7–9^

**Figure 1.**
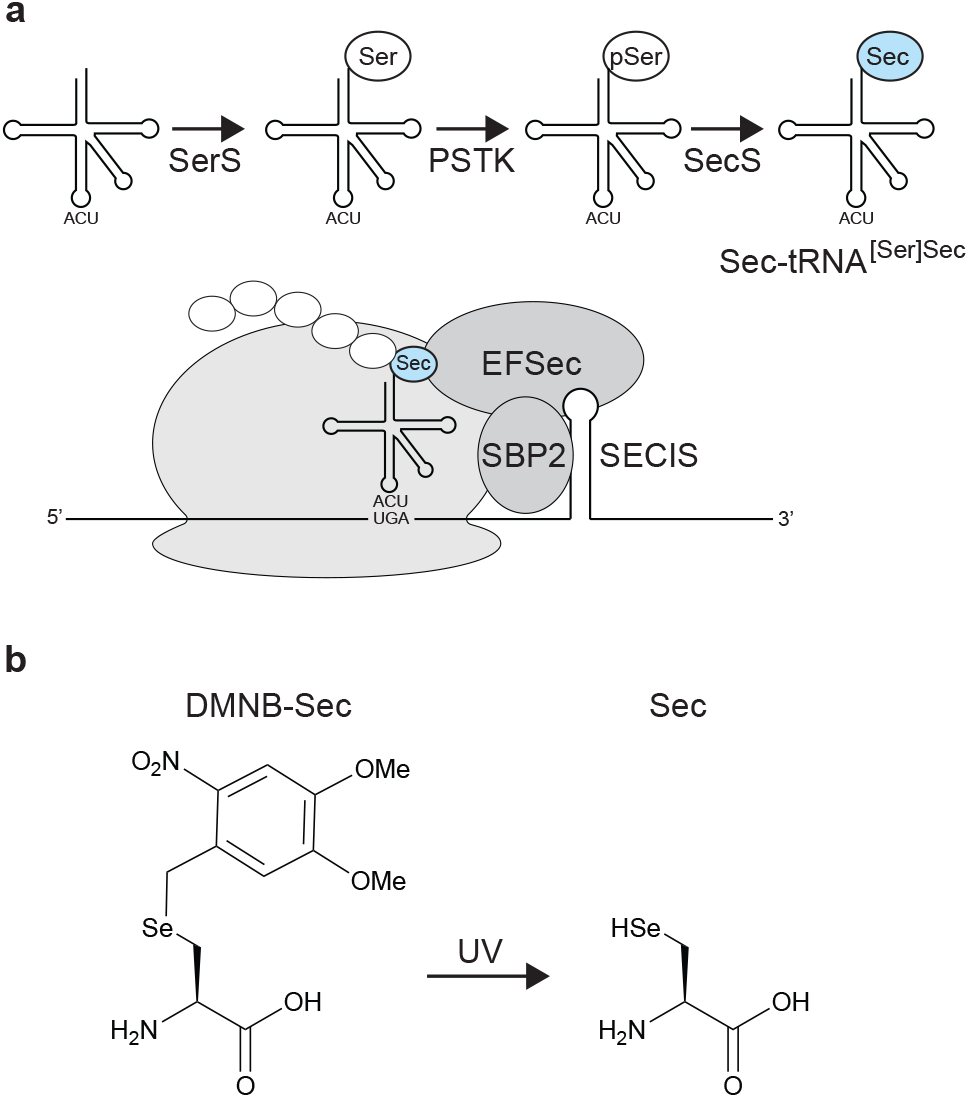
Selenocysteine incorporation and DMNB-Sec structure. (a) Endogenous eukaryotic Sec-incorporation mechanism. (b) Structures of DMNB-Sec and Sec.

Due to the complexity of endogenous selenoprotein production in eukaryotes, recombinant expression has proven challenging. The low efficiency of traditional manipulations, such as overexpression and mutation, has hindered the investigation of selenoproteins directly in mammalian cells. As a result, the biochemical and cell-biological functions of many human selenoproteins remain poorly characterized.^1^

Selenoproteins can be produced using expressed protein ligation, but is limited to *in vitro* studies, and necessitates refolding of the protein.^10^ A number of strategies have been employed to enhance the recombinant expression of selenoproteins in *E. coli*, including the use of reassigned sense codons, engineered tRNAs, and recoded release factor 1 deficient E. coli.^11–15^ While useful, these prokaryotic expression systems are frequently poorly suited for mammalian protein expression, and is not conducive to studying mammalian selenoproteins in their native cellular environment. In eukaryotic cells, unique SECIS elements from *Toxoplasma* have been used to improve selenoprotein production, but is dependent on Se availability.^16^

Genetic code expansion (GCE) technology allows site-specific incorporation of noncanonical amino acids using engineered aminoacyl-tRNA synthetase (RS)/tRNA pairs.^17–19^ GCE offers an attractive alternative for Sec incorporation that overrides many limitations of the endogenous pathway. Furthermore, protected analogs of Sec can be incorporated to produce selenoproteins that can be subsequently activated on demand. An allyl-protected Sec (ASec) and a photocaged (methyl-(6-nitropiperonyloxymethyl), MeNPOM) Sec (PSc) have been genetically encoded in *E. coli* using the pyrrolysyl-tRNA synthetase/tRNA pair, and can be uncaged to Sec using Pd-catalysis and light, respectively. ^20,21^ A second photo-caged (4,5-dimethoxy-2-nitrobenzyl, DMNB) Sec (DMNB-Sec) was incorporated in yeast using the *E. coli* leucyl-tRNA synthetase/tRNA pair. ^22^ Of these, only ASec has been successful applied to mammalian cells. ASec incorporation in mammalian cells used the engineered pyrrolysyl pair, but was restricted to a resilient model protein (EGFP); whether this incorporation system is sufficiently robust to express more sensitive endogenous selenoproteins is unclear. Furthermore, decaging ASec requires treatment with toxic palladium species, which severely limits its applicability in live mammalian cells. Palladium-mediated uncaging also affords poor temporal control, due to the time for diffusion into the cell, and the slow kinetics of the uncaging reaction. Finally, the necessary concentration of ASec in the culture media (0.2 mM) is toxic to mammalian cells, and necessitates supplementation with high concentration of cysteine (3.2 mM) to mitigate toxicity. ^21^

Here, we report a robust platform to incorporate photo-caged DMNB-Sec (Figure 1b) into native selenoproteins in mammalian cells. This platform presents distinct advantages, including: (1) generation of the native selenoprotein directly in mammalian cells, instead of in model organisms, such as *E. coli* or yeast; (2) production of a protected selenoprotein that is not susceptible to oxidative damage and inactivation prior to biochemical interrogation; (3) use of a minimally toxic DMNB-Sec amino acid at low concentrations (12.5-100 μM) without cysteine supplementation; and, (4) rapid uncaging by 365 nm irradiation, which is minimally invasive, and provides excellent spatial and temporal control.^23^

DMNB-Sec (Figure 1b) was synthesized as previously reported.^22^ Incorporation in mammalian cells was achieved using the *E. coli* LeuRS BH5 T252A/tRNA pair, which was originally evolved for photocaged serine (DMNB-Ser)^24^, and subsequently used for DMNB-Sec in yeast.^22^ LeuRS and 8 copies of the Leu tRNA were cloned into the pAcBac2 vector containing an eGFP-39-TAG reporter (Figure S1).^25–27^ Transient transfection into HEK293T cells was followed by growth in the presence or absence of 100 μM DMNB-Sec. Fluorescence of eGFP was 5-fold higher for lysates from cells exposed to DMNB-Sec (Figure 2a), consistent with the yeast LeuRS system with DMNB-Sec and DMNB-Cys.^22^

**Figure 2.**
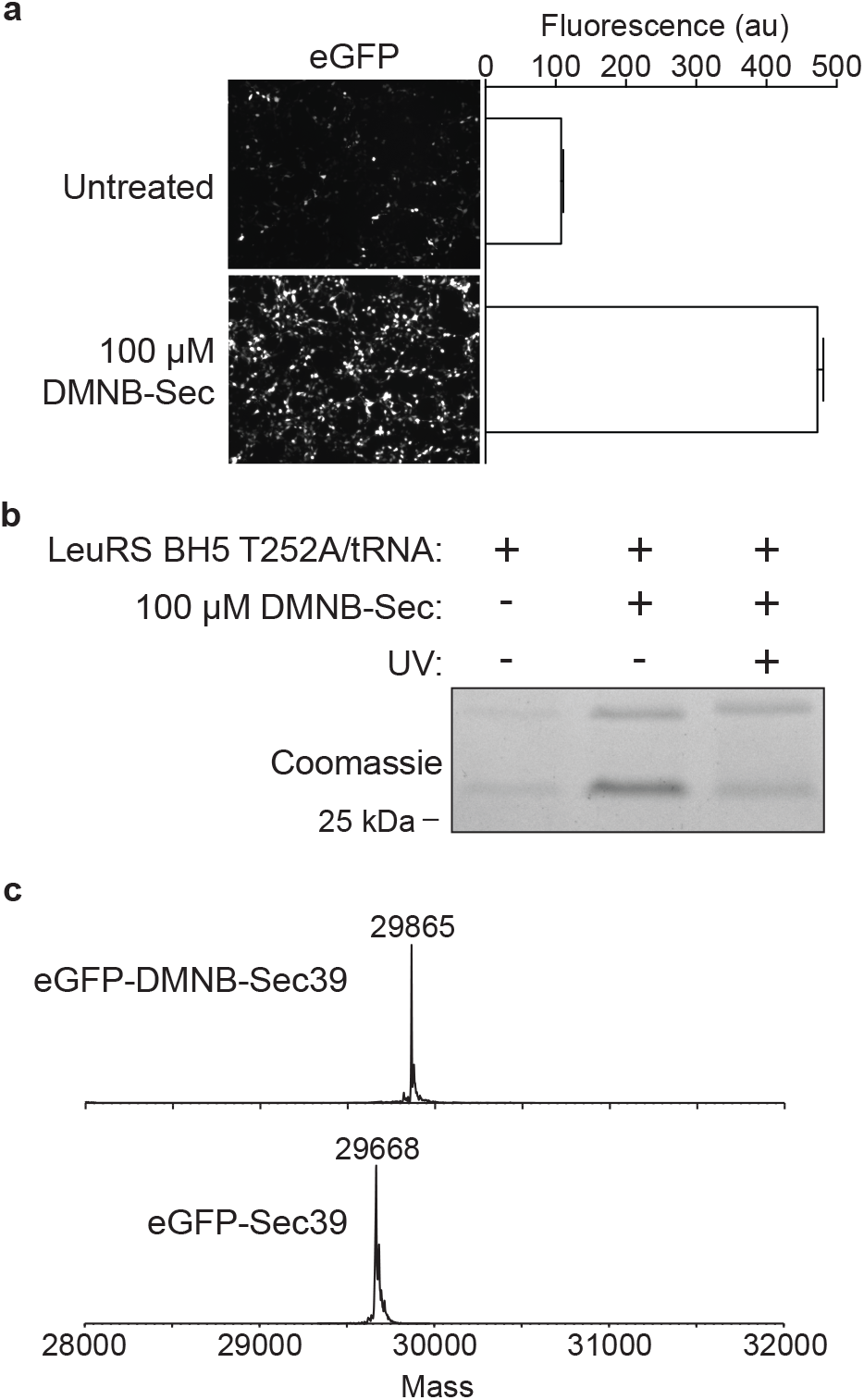
Generation of eGFP-DMNB-Sec39. (a) Fluorescence microscopy and quantification of HEK293T cells transiently transfected with a pAcBac2 plasmid encoding LeuRS BH5 T252A /Leu tRNA/eGFP-39-TAG, in the presence and absence of 100 μM DMNB-Sec. (b) SDS-PAGE of eGFP-DMNB-Sec39 purified from cells with and without UV irradiation. (c) ESI-MS analysis of eGFP-DMNB-Sec39 (expected mass: 29865.33) and eGFP-Sec39 (expected mass: 29669.24)

eGFP from non-irradiated cells, and cells grown in the absence of DMNB-Sec, were purified via a C-terminal 6XHis tag (Figure 2b). To determine uncaging efficiency, eGFP-DMNB-Sec39 expressing cells were irradiated (365 nm, 10 mins) (Figure 2b). Intact-protein electrospray-ionization mass spectrometry (ESI-MS) of these purified proteins confirmed the presence of the expected DMNB-Sec and Sec-containing eGFP species (Figures 2c, S2, S3). DMNB-Sec uncaging in yeast lysates resulted in the formation of diselenide dimers as well as dehydroalanine species.^22^ However, only the expected monomer was observed in ESI-MS analysis of mammalian-expressed eGFP-Sec39. Analysis of eGFP from cells grown in the absence of DMNB-Sec identified a mass consistent with the incorporation of leucine (Leu) or isoleucine (Ile) (Figure S4), which likely explains the low levels of eGFP fluorescence observed in the absence of DMNB-Sec (Figure 2a). When eGFP was expressed in the presence of 100 μM DMNB-Sec, eGFP-Leu/Ile39 was not detectable, suggesting that incorporation of these natural amino acids is suppressed by the presence of DMNB-Sec.

We then proceeded to utilize this platform to generate a photocaged version of the native mammalian selenoprotein, human methionine-*R*-sulfoxide reductase 1 (MsrB1). MsrB1 is localized to the cytoplasm and nucleus, and utilizes an active-site Sec to stereo-selectively reduce methionine-*R*-sulfoxide.^28^ Intriguingly, the MsrB1 homologs, MsrA, MsrB2 and MsrB3, utilize a Cys, instead of Sec, to perform similar chemistry. Studies comparing Cys-vs. Sec-containing Msr proteins have shown that Sec is advantageous for reductase activity.^1,29-31^ MsrB1 protects proteins from oxidative damage, and enables signaling via dynamic, enzyme-dependent methionine oxidation and reduction.^32,33^ The role of MsrB1 has been well characterized for the substrate actin, but regulation of other potential substrates is poorly understood, due in part to the difficulties with overexpression and temporally controlled activation of MsrB1 in mammalian cells. Expression of MsrB1 with a caged active-site Sec residue would facilitate further characterization of the cellular functions of MsrB1.

To recombinantly express MsrB1, a pAcBac2 plasmid containing the following elements was generated: (1) LeuRS BH5 T252A; (2) 8 copies of the *E. coli* Leu tRNA; and, (3) the MsrB1 gene with the amber stop codon (MsrB1-TAG95) or a Cys codon (MsrB1-Cys95) at residue 95 (the site of the endogenous Sec residue), and a C-terminal 6XHis tag. To test for successful incorporation of DMNB-Sec, MsrB1 expression was monitored via immunoblot using antibodies against MsrB1 and the C-terminal 6XHis tag. Importantly, the 6XHis antibody will only detect full-length protein generated via successful amber suppression. Full-length MsrB1 protein was observed for MsrB1-Cys95, and MsrB1-TAG95 cells only in the presence of DMNB-Sec (Figure 3a). As expected, no protein was detected in the MsrB1-TAG95 cells in the absence of DMNB-Sec.

**Figure 3.**
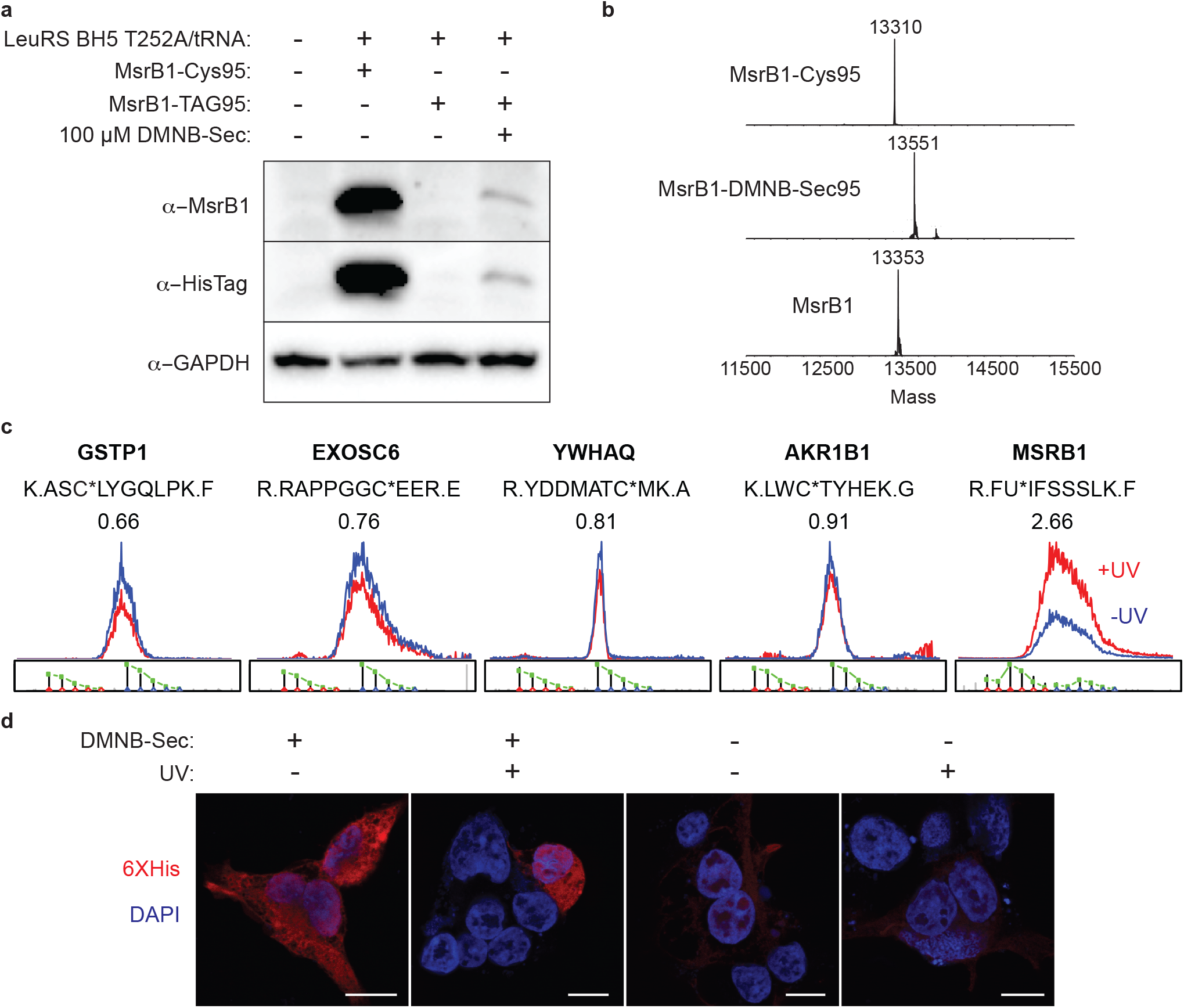
Generation of MsrB1-DMNBSec95. (a) Immunoblot of MsrB1 expression with a Cys or TAG amber codon at residue 95 in the presence and absence of 100 μM DMNB-Sec. (b) ESI-MS analysis of MsrB1-Cys95 (expected mass: 13309.97), MsrB1-DMNB-Sec95 (expected mass: 13552.04), and uncaged MsrB1-Sec95 (expected mass: 13356.87). (c) Low pH isoTOP-ABPP analysis showing extracted ion chromatograms for Cys- or Sec-containing peptides from irradiated (red, IA-light) or non-irradiated (blue, IA-heavy) cells expressing MsrB1-DMNBSec95. (d) Immunofluorescence of cells transfected with MsrB1-TAG95 in the presence or absence of DMNB-Sec, before and after irradiation. Scale bar represents 10 μm.

Purification and analysis by ESI-MS confirmed the presence of MsrB1-Cys95 and MsrB1-DMNBSec95 (Figures 3b, S5, S6). Successful uncaging was confirmed by subjecting purified MsrB1-DMNBSec95 to UV irradiation (365 nm, 10 mins), and subsequent ESI-MS analysis (Figure 3b, S7). As with eGFP-Sec39, dimer and dehydroalanine formation was not observed by ESI-MS. Additionally, MsrB1-Ile/Leu39 was not detected, confirming that misincorporation of Ile/Leu is negligible in the presence of DMNB-Sec.

To demonstrate uncaging and activation of MsrB1 directly in cells, we utilized an isoTOP-ABPP method that was modified for selenoprotein detection.^34^ Typical isoTOP-ABPP monitors Cys reactivity in complex proteomes via the following steps: (1) cell-lysate labeling with an iodoacetamide (IA)-alkyne probe; (2) conjugation of IA-alkyne modified proteins to a cleavable-biotin tag using copper (I)-catalyzed azide-alkyne cycloaddition (CuAAC); (3) enrichment of IA-alkyne modified proteins on streptavidin beads; (4) on-bead trypsin digestion and cleavage of the linker to release IA-alkyne modified peptides for analysis by LC/LC-MS/MS. Incorporation of isotopically light and heavy IA-alkyne probes, or cleavable linkers, enable quantitative comparisons of cysteine reactivity across different biological samples.^35,36,37^ Performing the initial IA-alkyne labeling step at low pH (~pH 5.75) was shown to suppress Cys labeling, and thereby enhance the detection of Sec-containing peptides for the analysis of Sec-reactivity changes.^34^

Low-pH isoTOP-ABPP analysis was used to compare lysates from irradiated (IA-light labeled) and non-irradiated (IA-heavy labeled) HEK cells expressing MsrB1-DMNB-Sec95 (Figure S8). The IA-labeled MsrB1 Sec-containing peptide (R.FU*IFSSSLK.F) was identified via LC/LC-MS/MS, and shown to display a light:heavy ratio of 2.66, indicative of ~3-fold increased Sec reactivity in the irradiated sample (Figure 3c). In contrast, identified Cys-containing peptides showed R values of ~1 (Figure 3c), indicating no reactivity change upon irradiation. Importantly, the identified MsrB1 peptide displayed an isotopic envelope characteristic of Se-containing peptide species, confirming Sec incorporation.^38^ Together, the data from the low pH isoTOP-ABPP analysis confirms the successful incorporation and uncaging of DMNB-Sec in MsrB1 directly in mammalian cells.

Endogenous MsrB1 is known to localize to the cytosol and nucleus of cells.^28^ To confirm that MsrB1-DMNB-Sec95 shows similar localization patterns, we utilized confocal microscopy to determine the cellular localization of MsrB1-DMNB-Sec95 in irradiated and non-irradiated cells. Regardless of irradiation, the recombinant MsrB1 was shown to display both cytosolic and nuclear localization, consistent with the endogenous protein (Figure 3d). As expected, cells cultured in the absence of DMNB-Sec did not show a significant MsrB1 signal. These imaging studies confirm that photocaged versions of endogenous selenoproteins can be produced and activated in mammalian cells with the expected cellular localization for subsequent biological interrogation.

In summary, studies of selenoprotein function have been hindered by the poor efficiency of recombinant expression of Sec-containing proteins and related mutants. Furthermore, the susceptibility of Sec residues to oxidation results in rapid oxidative inactivation of proteins, limiting the ability to study the active form of the protein in cells. Here, a photocaged DMNB-Sec is incorporated into proteins in mammalian cells through genetic code expansion. The corresponding Sec-containing protein can be generated by UV irradiation, which is compatible with live cells, and is minimally disruptive to protein function. We report the first incorporation of a protected Sec amino acid into a native selenoprotein directly in mammalian cells, with temporal control of selenoprotein activity. Of note, the all-in-one pAcBac2 vector system used to deliver the DMNB-Sec incorporation machinery can be readily packaged into an engineered baculovirus vector that can facilitate highly efficient unnatural amino acid incorporation into a wide variety of mammalian cells and tissues, including challenging cells such as neurons and stem cells.^25,27^ Efforts are also underway to improve the efficiency of DMNB-Sec incorporation through optimization of the LeuRS/tRNA pair. The technology described herein lays the foundation for expressing temporally controlled selenoproteins in mammalian cells for further advancing our understanding of selenoprotein biology.

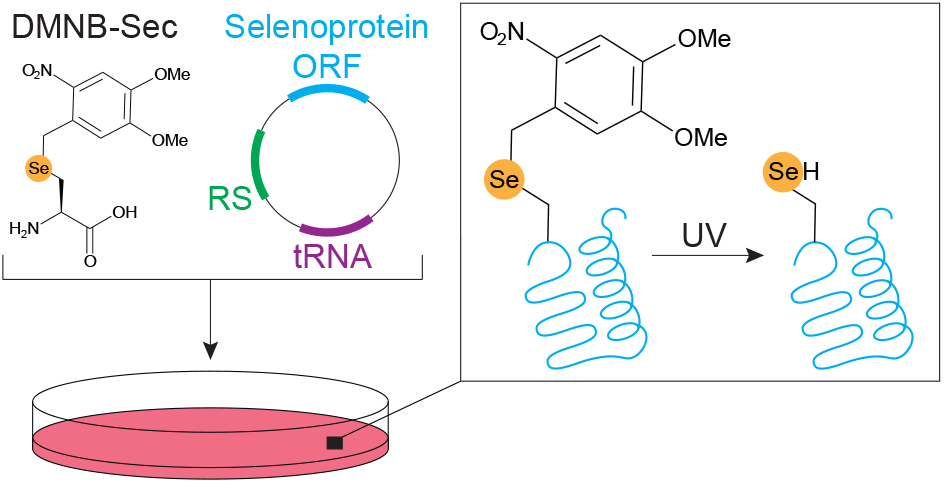

## ASSOCIATED CONTENT

### Supporting Information

Supporting information includes a methods section and supporting figures.

## ACKNOWLEDGMENT

We thank Bret Judson and the Boston College Imaging Core for infrastructure and support, and members of the We-erapana and Chatterjee Labs for helpful discussions.

