## Supporting information for "Generation of Recombinant Mammalian Selenoproteins through Genetic Code Expansion with Photocaged Selenocysteine"

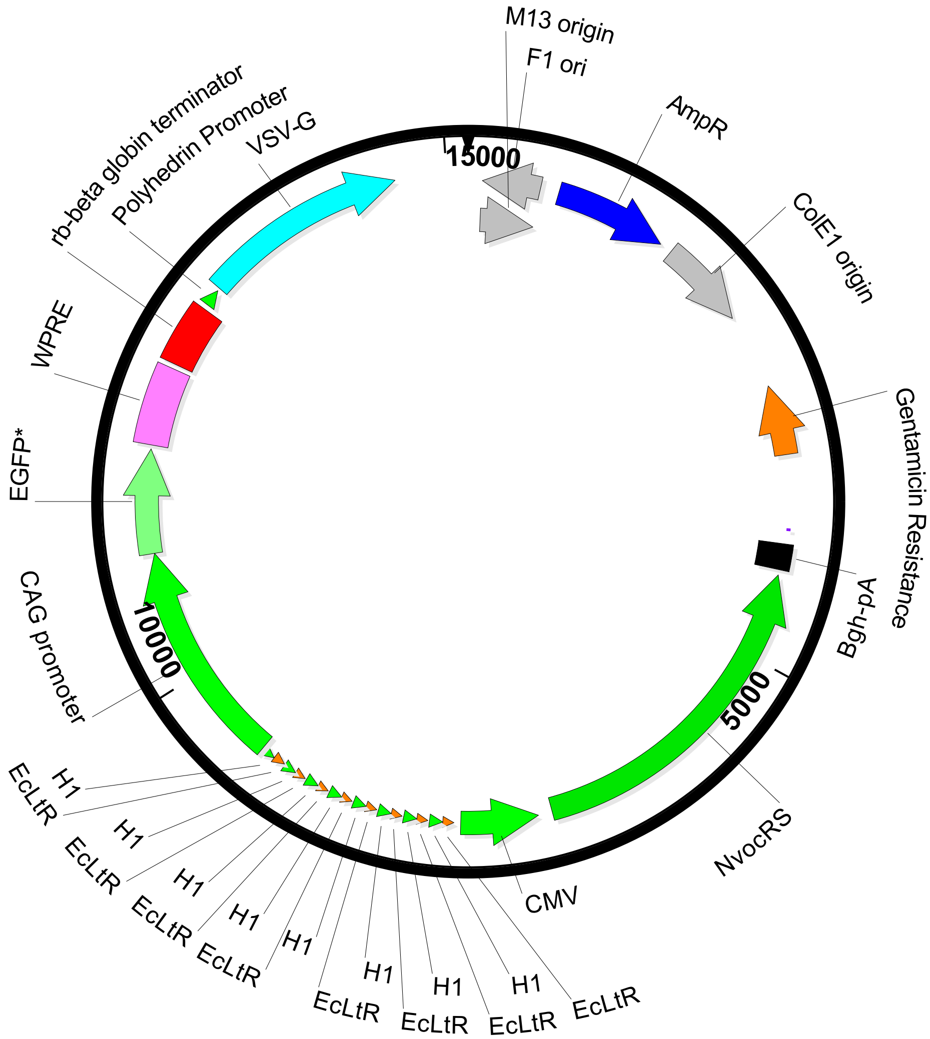


**Supplemental Figure 1.** Plasmid map of pAcBac2- EcLeuRS-BH5 T252A plasmid

**a**

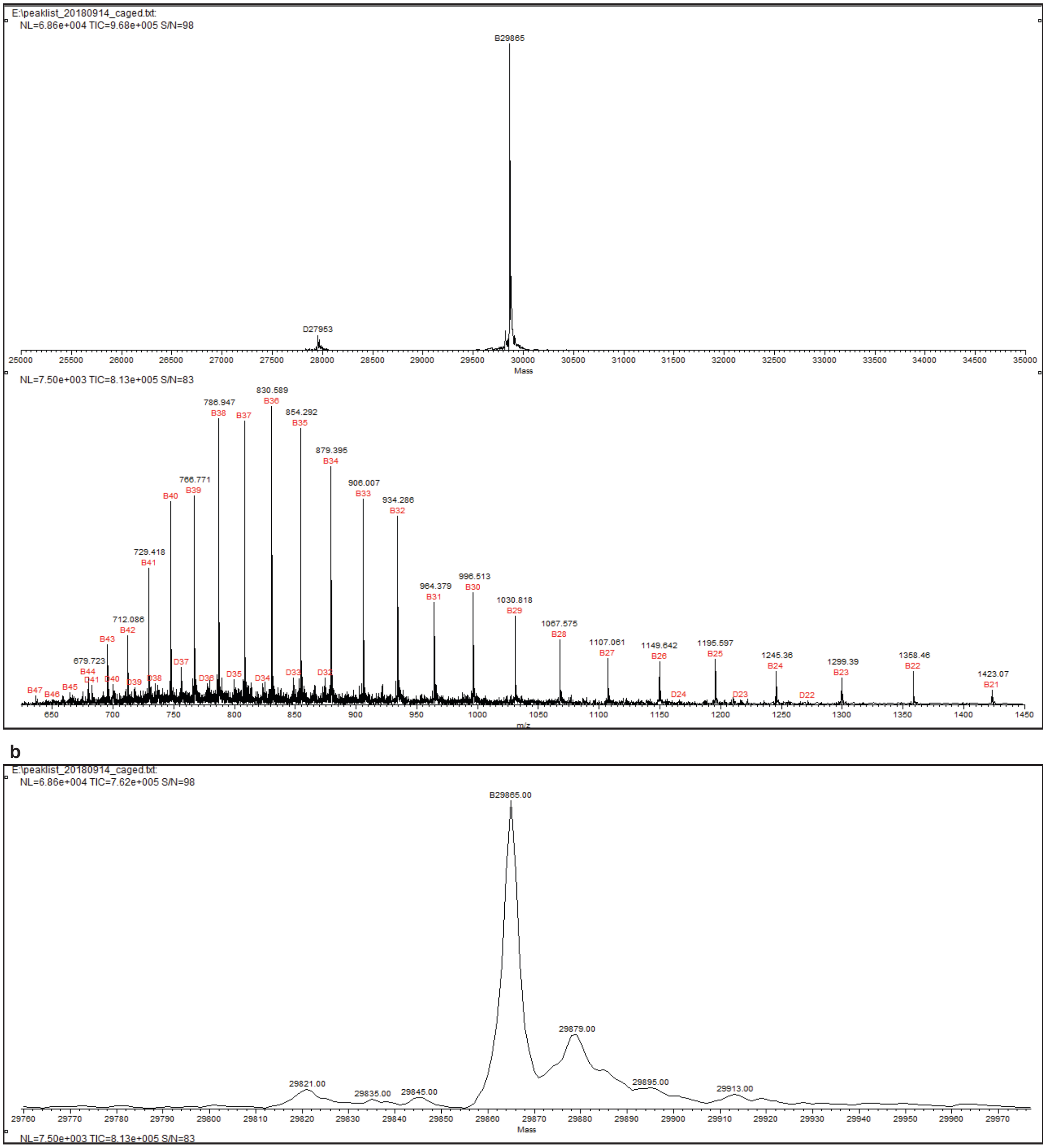


**Supplemental Figure 2.** ESI-MS of eGFP-DMNBSec39 (a) Deconvoluted peak shown with mass range from 25000 to 35000 m/z (expected: 29865.33) and spectrum (b) Zoomed in view of deconvoluted peak at 29865 m/z.

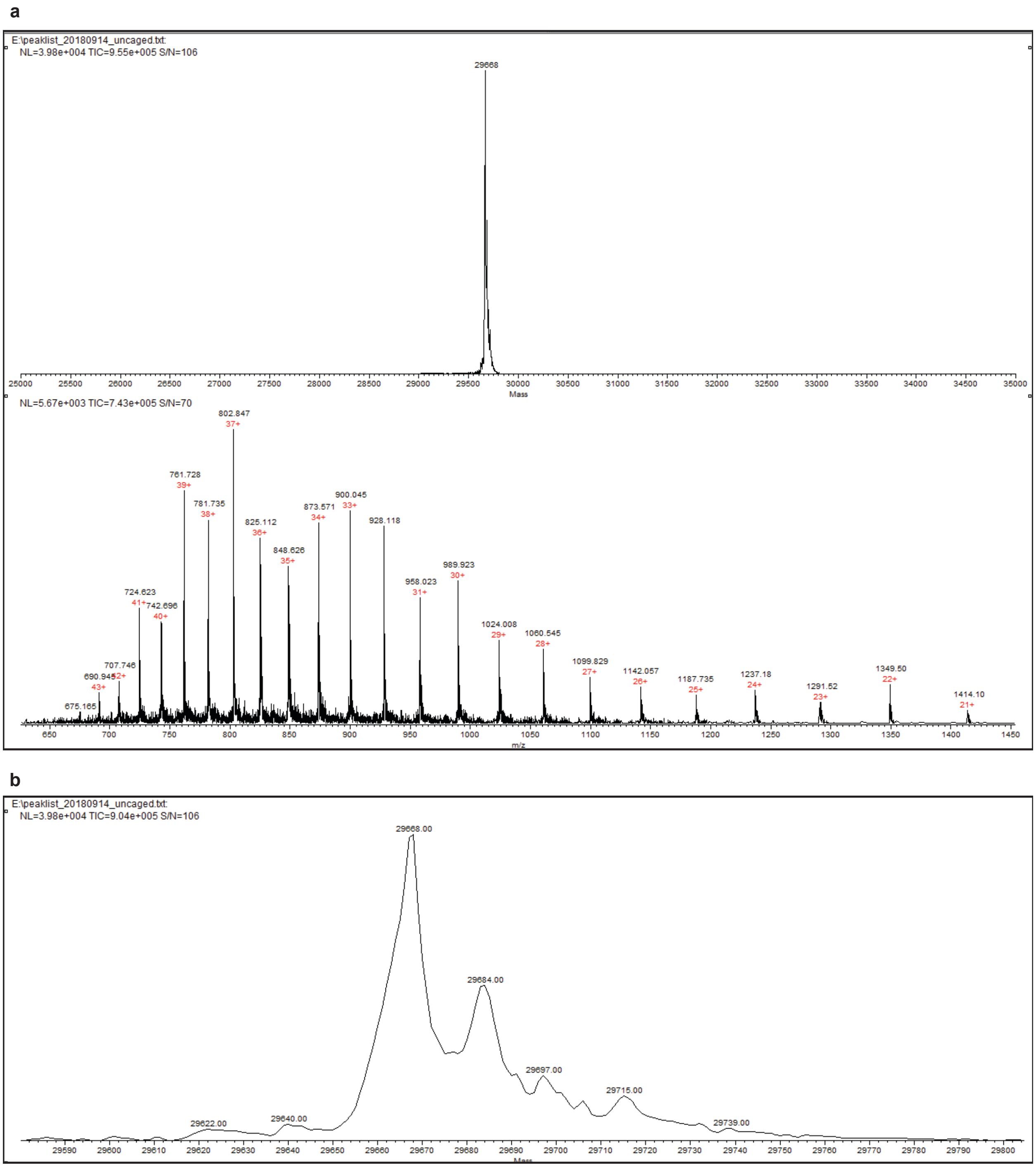


**Supplemental Figure 3.** ESI-MS of eGFP-Sec39 (a) Deconvoluted peak shown with mass range from 25000 to 35000 m/z (expected: 29669.24) and spectrum (b) Zoomed in view of deconvoluted peak at 29668 m/z.

a

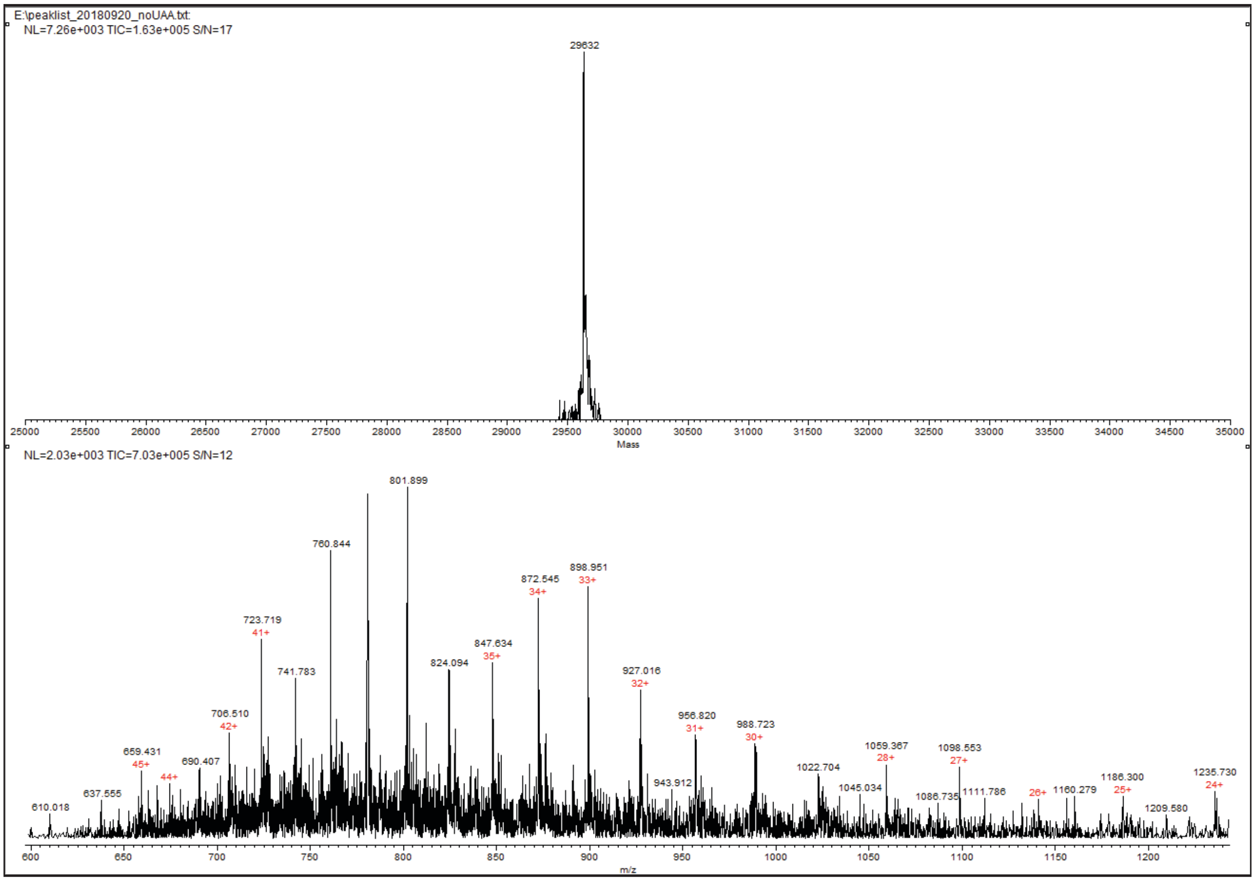


b

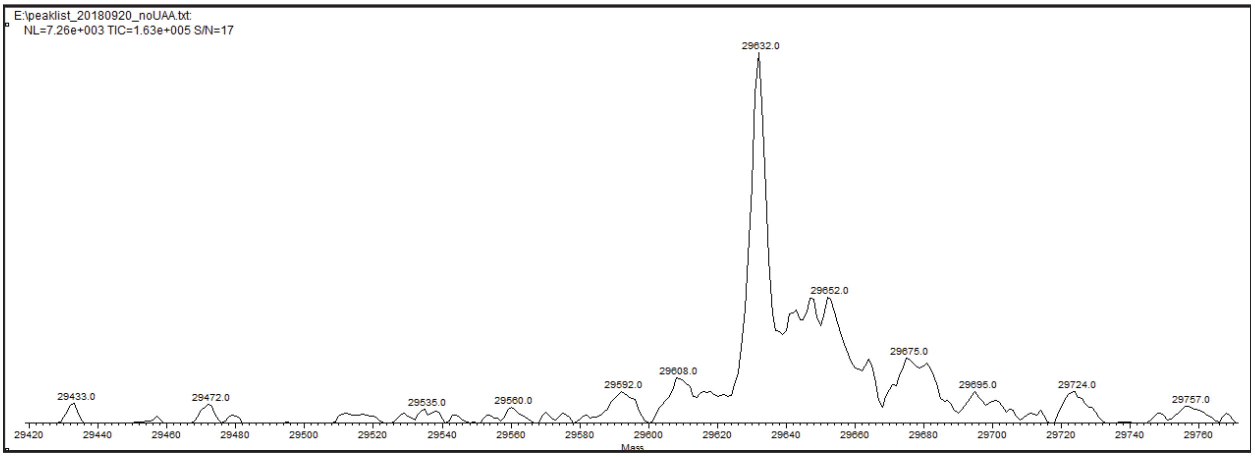


**Supplemental Figure 4.** ESI-MS of eGFP-TAG-39 grown in the absence of DMNB-Sec (a) Deconvoluted peak shown with mass range from 25000 to 35000 m/z and spectrum. The expected mass for eGFP with Leu/Ile incorporated at position 39 is 29632.350. Leu incorporation is likely due to retained ability of the engineered *E. coli* Leu RS to charge tRNA with Leu in the absence of a preferred substrate. (b) Zoomed in view of deconvoluted peak at 29632 m/z.

**a**

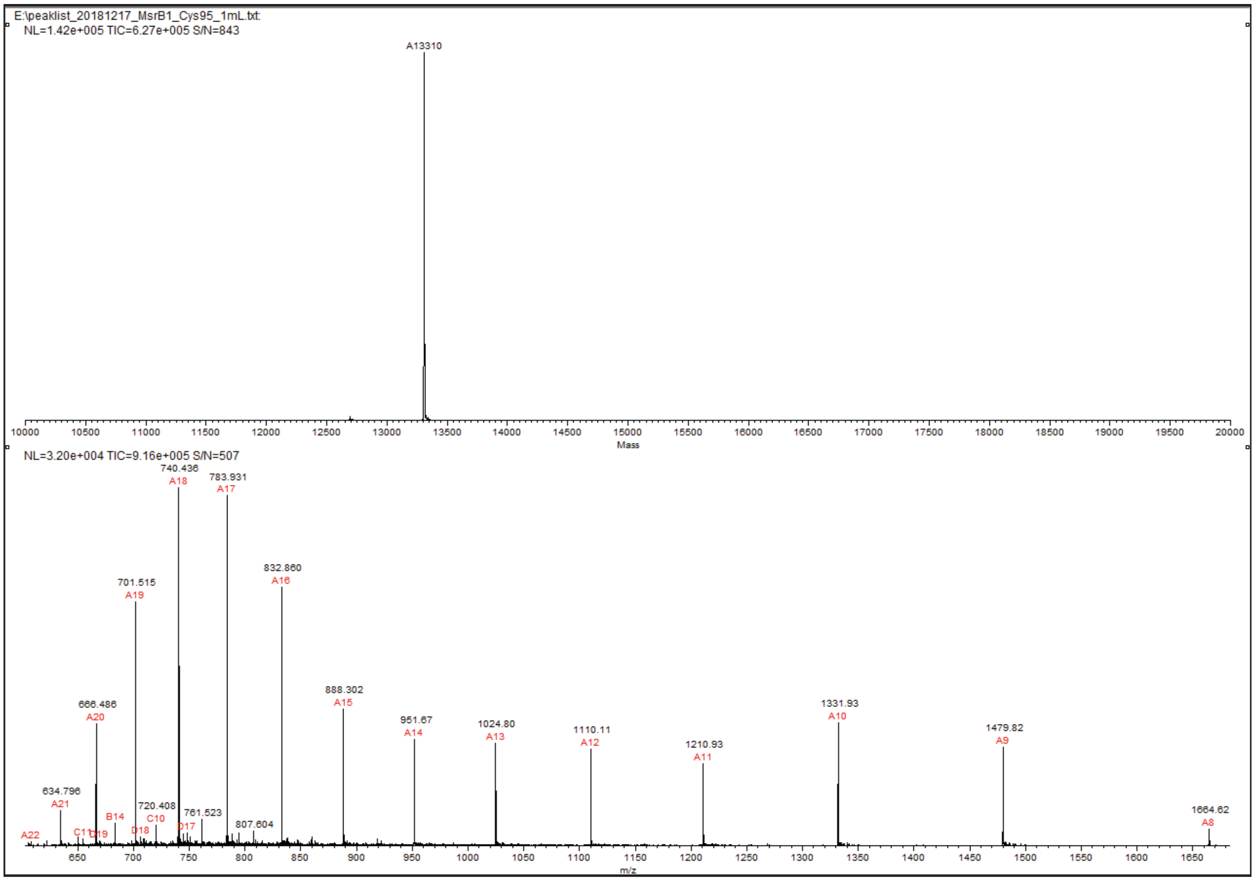


**b**

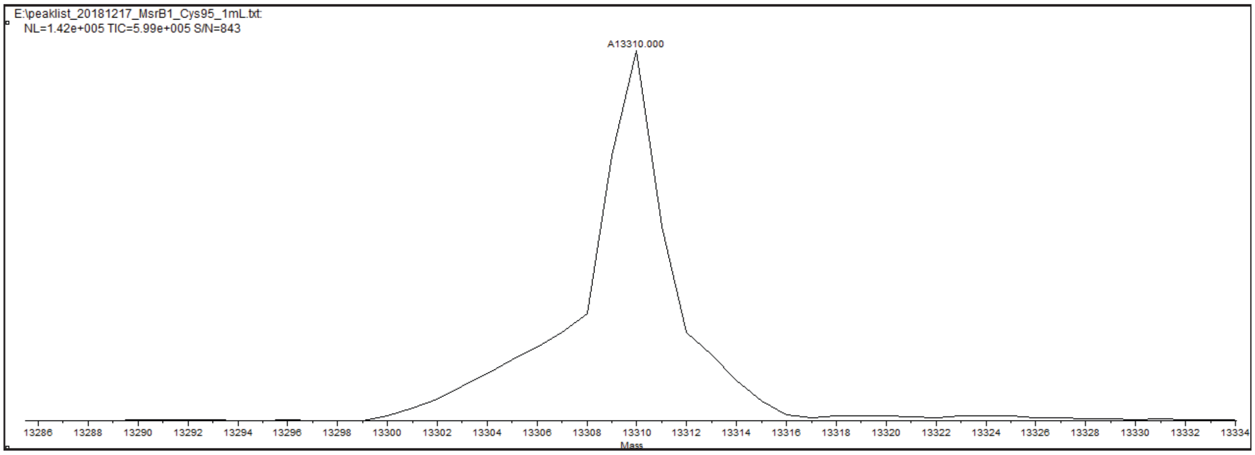


**Supplemental Figure 5.** ESI-MS of MsrB1-Cys95 (a) Deconvoluted peak shown with mass range from 10000 to 20000 m/z (expected: 13309.97) and spectrum (b) Zoomed in view of deconvoluted peak at 13310 m/z.

a

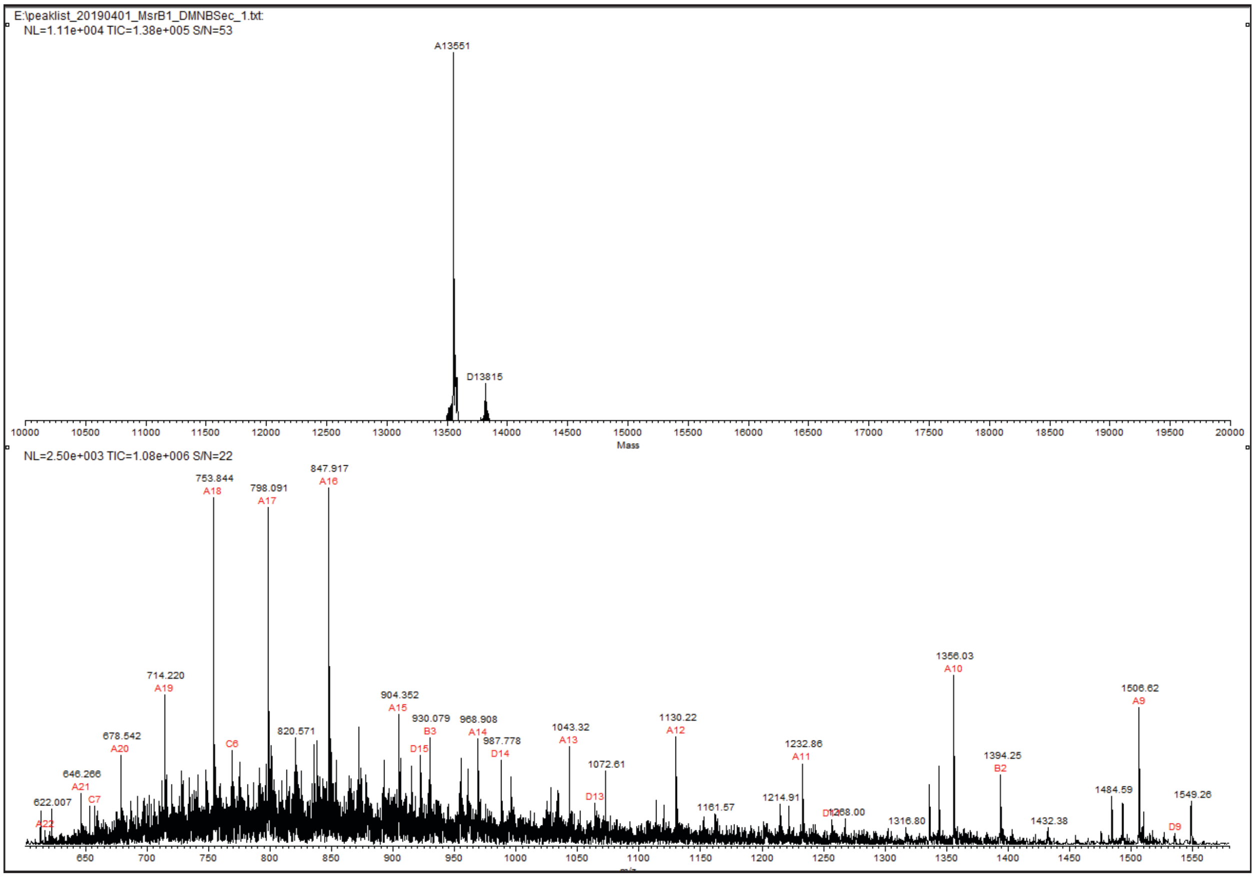


b

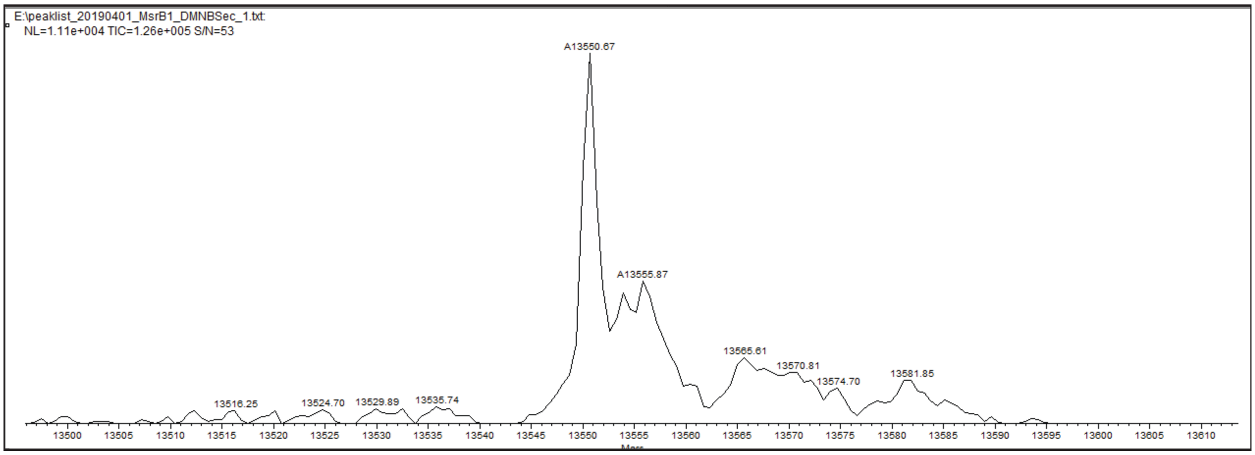


**Supplemental Figure 6.** ESI-MS of MsrB1-DMNB-Sec95 (a) Deconvoluted peak shown with mass range from 10000 to 20000 m/z (expected: 13552.04) and spectrum (b) Zoomed in view of deconvoluted peak at 13551 m/z.

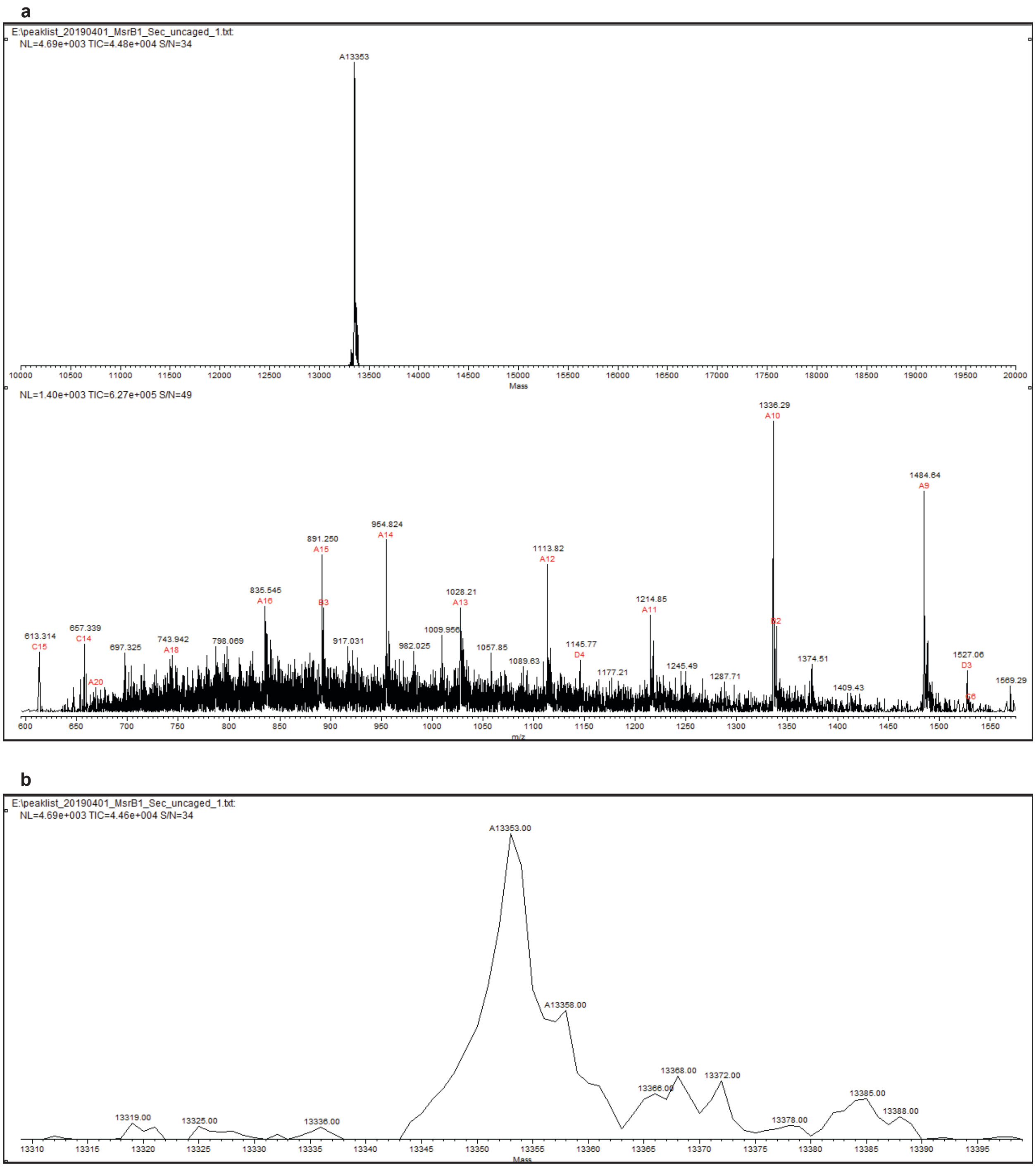


**Supplemental Figure 7.** ESI-MS of MsrB1-Sec95 (a) Deconvoluted peak shown with mass range from 10000 to 20000 m/z (expected: 13356.87) and spectrum (b) Zoomed in view of deconvoluted peak at 13553 m/z.

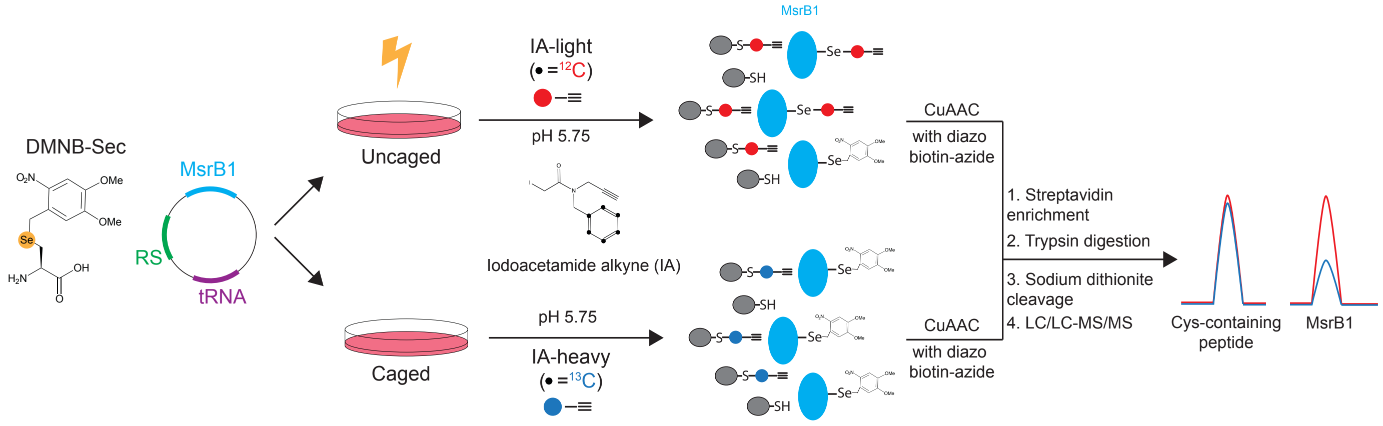


**Supplemental Figure 8**. Selenoprotein peptide enrichment strategy

### Methods

#### Synthesis of DMNB-Sec

DMNB-Sec was synthesized as previously reported. [1]

#### Cloning of pAcBac2- EcLeuRS-BH5 T252A plasmids

The eGFP-containing plasmid was generated from the previously reported pAcBac2R-Anap plasmid by replacing the AnapRS with the EcLeuRS-BH5 T252A mutant.[2] The MsrB1-containing plasmid was subsequently generated by replacing the eGFP ORF with the ORF for human MsrB1-6XHis with either cysteine or the amber codon TAG at residue 95, the site of the native selenocysteine.

#### Expression of GFP-Sec and MsrB1

GFP-6XHis and MsrB1-6XHis variants were expressed in HEK293T cells. For GFP microscopy and fluorescence quantification, transfections were performed in 12-well tissue culture plates. Cells were transiently transfected using 21 μL of serum-free DMEM, 1 μg of plasmid DNA, and 4 μL of 1 mg/mL PEI MAX (Polysciences, 24765-1) and grown in the presence or absence of 100 μM DMNB-Sec for 48 hours.

For protein purification and proteomic analysis, GFP-6XHis variants and MsrB1-6XHis variants were expressed in 10 cm tissue culture dishes, using 210 μL of serum-free DMEM, 12 μg plasmid DNA, and 50 μL of 1 mg/mL PEI MAX for transfection. GFP-DMNBSec39-6XHis expression was performed in the presence of 100 μM DMNB-Sec and MsrB1-DMNBSec95-6XHis expression was performed in the presence of 12.5 μM DMNB-Sec.

In order to uncage GFP-DMNBSec39-6XHis and MsrB1-DMNBSec95-6xHis (generating GFP-Sec39-6XHis and MsrB1-6XHis), cells or purified proteins were irradiated at 365 nm for 10 min on ice using 120 W LED-array (Larson Electronics).

#### GFP quantification

Cells in a 12-well plate were solubilized in 200 μL of CelLyticM (Sigma Aldrich, C2978) and after a 15 min incubation at room temperature, 180 μL of lysate was moved to a 96 well plate. Lysates were measured for end point fluorescence with excitation at 488 nm, emission measured at 532 nm, a cutoff at 530 nm, and 100 reads per well using a SpectraMAX M5 (Molecular Devices). Fluorescence was corrected by subtracting the fluorescence of untransfected cells.

#### Protein Purification and LC-MS analysis

HEK293T cell pellets were solubilized in CelLyticM supplemented with Pierce Universal Nuclease (Thermo Scientific, 88700) and Halt Protease Inhibitor Cocktail

(Thermo Scientific, 87786). After 10 min of incubation, lysates were spun at 16800 g. GFP-6XHis and MsrB1-6XHis variants were purified using HisPur Ni-NTA Resin (Fisher Scientific, PI88221) following manufacturer's instructions. After purification, proteins were analyzed by SDS-PAGE and whole protein mass spectrometry (ESI-MS). ESI-MS was performed using a 1260 Agilent Infinity Series HPLC coupled with a 6230 Agilent TOF Mass Spectrometer. Anticipated masses for whole proteins were calculated to include the loss of initiator methionine and addition of acetylation.

**MsrB1 western blot**

HEK293T lysates were prepared in CelLyticM as described above. Lysates were normalized for protein concentration and run on a 15% polyacrylamide gel at 150 V. Proteins were transferred to a nitrocellulose membrane (Fisher Scientific, 45-004-003) at 75 V for 90 min. Membranes were blocked with 5% BSA in TBST for 1 hour at room temp, and then incubated with antibodies against MsrB1 (Santa Cruz Biotechnology, SC-34846), the 6X His epitope (Cell Signaling Technology, 2365), or GAPDH (Cell Signaling Technology, 2118) overnight at 4 °C. Membranes were washed 3 times with TBST and incubated with HRP-linked secondary antibodies (Abcam, ab97105 for the MsrB1 primary antibody, and Cell Signaling Technologies 7074 for His tag and GAPDH primary antibodies) for 1 hour at room temperature. Membranes were washed 3 times with TBST and incubated with SuperSignal West Pico PLUS chemiluminescent substrate (Fisher Scientific, PI34578). The membranes were imaged with the ChemiDoc MP imaging system (BioRad).

**Proteomics**

Pellets of HEK293T cells expressing either MsrB1-DMNBSec95-6XHis (caged) or MsrB1-6XHis (uncaged) were resuspended in Selenoprotein Enrichment (SE) buffer (50 mM MES, 50 mM $NaH_2PO_4$, 100 mM NaCl, pH 5.75). [3] For the uncaged pellet, the SE buffer contained 100 μM of iodoacetamide alkyne light (IA-light) while the caged pellet the SE buffer contained 100 μM of iodoacetamide alkyne heavy (IA-heavy). Pellets were sonicated for 3 rounds of 10 pulses at 75% amplitude. The lysates were spun down for 5 min at 10,000 g. The lysates were incubated for 1 hour at room temperature, and their protein concentration was determined using the DC Protein Assay (BioRad). Lysates (2.4 mg/mL) were then conjugated to the diazo biotin azide tag (Click Chemistry Tools, 1041-25) via CuAAC (100 μM of diazo biotin azide, 1 mM TCEP [Aldrich, C4706], 100 mM TBTA [Aldrich, 678937], and 1 mM CuSO4) for 1 hour at room temperature. Proteins were precipitated by centrifugation (6,500 g, 10 min, room temperature), and washed 3 times with ice-cold methanol (centrifuging at 6,500 g, 10 min, 4 °C). The protein pellet was then resuspended in 1 mL of 1.2% SDS in PBS by sonication and heating (5 min, 95 °C). Samples were diluted in 5 mL of PBS and incubated with 100 μL of streptavidin-agarose beads (Thermo Scientific, 20353) and rotated at 4 °C overnight. The beads were then incubated with rotation at room temperature for 3 hours, washed with 0.2% SDS/PBS (5 mL), PBS (3 x 5 mL), and water (3 x 5 mL). Between washes the beads were pelleted by centrifugation (1400 g, 3 min). The beads were transferred to screw-cap Eppendorf tubes and resuspended in 500 μL of 6 M Urea in PBS. Samples were treated with 10 mM

Ultrapure DTT (Invitrogen, 15508-013) and heated for 20 min at 65 °C. Samples were then alkylated with 20 mM iodoacetamide (ACROS, 122270050) and incubated at 37 °C for 30 min. The beads were pelleted by centrifugation and resuspended in 200 μL of 2 M Urea in PBS, 1 mM $CaCl_2$, and 2 mg sequence grade modified trypsin (Promega, V5111). The tryptic digestion was allowed to incubate overnight at 37 °C. The beads were washed in PBS (3 x 500 μL) and water (3 x 500 μL). Labeled peptides were eluted from the beads by sodium dithionite-mediated cleavage of the diazo biotin azide tag. Beads were incubated with 50 μL of 25 mM sodium dithionite (Sigma-Aldrich, 161527) in PBS, rotating at room temperature for 1 hour. After centrifugation, the supernatant was collected and saved. The beads were then incubated with 75 μL of 25 mM sodium dithionite at room temperature for 1 hour and finally 75 μL of 50 mM sodium dithionite at room temperature for 1 hour. All of the collected elutions were combined. The beads were washed twice more with 75 μL water, and the washes were combined with the elutions. Formic acid (17.5 μL, Sigma) was added to the samples (350 μL), and the samples were stored at -20 °C.

Mass spectrometry was performed using a Thermo LTQ Orbitrap Discovery mass spectrometer coupled to an Agilent 1200 series HPLC. Labeled peptide samples were pressure loaded onto 250 mm fused silica desalting column packed with 4 cm of Aqua C18 reverse phase resin (Phenomenex). Peptides were eluted onto a 100 mm fused silica biphasic column packed with 10 cm C18 resin and 4 cm Partisphere strong cation exchange resin (SCX, Whatman), using a five-step multidimensional LC-MS protocol (MudPIT). Each of the five steps used a salt push (0%, 50%, 80%, 100%, and 100%), followed by a gradient of 5-100% buffer B in Buffer A (Buffer A: 95% water, 5% acetonitrile, 0.1% formic acid; Buffer B: 20% water, 80% acetonitrile, 0.1% formic acid). The flow rate through the column was approximately 0.25 mL/min, with a spray voltage of 2.75 kV.

In order to detect MsrB1 peptides, a mass list was used to target the +2 and +3 ions for the Sec-containing peptide in MsrB1 (masses were 721.83 and 481.56 respectively, including IA-light and the cleaved diazo biotin azide tag). One full MS1 scan (400-1800 MW) was followed by 2 data-dependent scans and dynamic exclusion was disabled.

The tandem MS data, generated from the 5 MudPIT runs, was analyzed by the SEQUEST algorithm. The precursor-ion mass tolerance was set at 50 ppm while the fragment-ion mass tolerance was set to 0 (default setting). Static modification of Cys residues (+57.0215, iodoacetamide alkylation) was assumed with no enzyme specificity. The datasets were searched against human reverse-concatenated non-redundant UniProt Database (release-2012_11). These datasets were searched for differential Cys modification with IA-light (+306.1481) or IA-heavy (+312.1682) with cleaved diazo biotin azide. Considering SEQUEST does not designate a Sec residue, the datasets collected were searched against a modified UniProt database where all Sec residues were replaced with Cys. These searches were performed with differential modifications on Cys that account for the increased mass of Sec as well as IA-light or IA-heavy and cleaved diazo biotin azide (+354.0925 and +360.1126).

MS2 spectra matches were assembled into protein identifications and filtered using DTASelect2.0 to generate a list of protein hits with a peptide false-discovery rate of 5%, with the –trypstat and –modstat options applied. Peptides were restricted to fully tryptic (-

y 2) with a found modification (-m 0) and a delta-CN score greater than 0.06 (-d 0.06). Single peptides per locus were also allowed (-p 1) as were redundant peptides identifications from multiple proteins, but the database contained only a single consensus splice variant for each protein.

Peptide light to heavy (L:H) ratios were calculated using the Cimage quantification package. [4] In order to quantify L:H ratios from MsrB1 peptides residue an additional column and row for selenium and selenocysteine was included into the cimage.params table and modified Cys residues in the DTASelect_filter.txt file were edited back to Sec.

**Immunofluorescence**

HEK293T cells were transfected as above and treated with or without 12.5 μM DMNBSec. After 24 hours cells were trypsinized and split into 35 mm glassbottom dishes (MatTek Corporation, P35G-1.5-10-C) coated with poly-D-lysine (Sigma-Aldrich, P6407-5MG). After a further 24 hours, cells were irradiated (365 nm, 10 min), returned to the incubator for 10 minutes, and then fixed with 4% paraformaldehyde in PBS (diluted from Thermo PI28906) for 15 minutes at room temperature. Plates were washed 5 times 5 minutes with PBS and blocked for 1 hour at room temperature in PBS with 5% normal serum and 0.3% Tween-20. Cells were incubated with an antibody raised against the C-term 6XHis tag (Cell Signaling Technologies, 12698) at a 1:400 dilution PBS with 1% BSA and 0.3% Tween-20 overnight at 4 °C. Cells were washed 3 times 5 minutes with PBS, then incubated with an anti-rabbit-A594 secondary antibody (Cell Signaling Technologies, 8889) diluted 1:500 in PBS with 1% BSA and 0.3% Tween-20 for 1 hour at room temperature. Cells were washed 3 times 5 minutes, and then mounted with Prolong Gold Antifade reagent with DAPI (Cell Signaling Technologies, 8961S). Images were acquired using a Leica (Wetzlar, Germany) TCS SP5 scanning confocal microscope using a Plan-Apochromat 63×/1.40 numerical aperture lens. Images were exported from Leica Application Suite Advanced Fluorescence (LAS AF) software as .tiffs and processed in Adobe Photoshop CC 2019.
